# Parallel cognitive maps for short-term statistical and long-term semantic relationships in the hippocampal formation

**DOI:** 10.1101/2022.08.29.505742

**Authors:** Xiaochen Y. Zheng, Martin N. Hebart, Raymond J. Dolan, Christian F. Doeller, Roshan Cools, Mona M. Garvert

**Author notes:** Correspondence to Xiaochen Y. Zheng, Donders Centre for Cognitive Neuroimaging, Kapittelweg 29, 6525 EN, Nijmegen, The Netherlands.

## Abstract

The hippocampal-entorhinal system uses cognitive maps to represent spatial knowledge and other types of relational information, such as the transition probabilities between objects. However, objects can often be characterized in terms of different types of relations simultaneously, e.g. semantic similarities learned over the course of a lifetime as well as transitions experienced over a brief timeframe in an experimental setting. Here we ask how the hippocampal formation handles the embedding of stimuli in multiple relational structures that differ vastly in terms of their mode and timescale of acquisition: Does it integrate the different stimulus dimensions into one conjunctive map, or is each dimension represented in a parallel map? To this end, we reanalyzed functional magnetic resonance imaging (fMRI) data from Garvert et al. (2017) that had previously revealed an entorhinal map which coded for newly learnt statistical regularities. We used a triplet odd-one-out task to construct a semantic distance matrix for presented items and applied fMRI adaptation analysis to show that the degree of similarity of representations in bilateral hippocampus decreases as a function of semantic distance between presented objects. Importantly, while both maps localize to the hippocampal formation, this semantic map is anatomically distinct from the originally described entorhinal map. This finding supports the idea that the hippocampal-entorhinal system forms parallel cognitive maps reflecting the embedding of objects in diverse relational structures.

## Introduction

The hippocampal-entorhinal system builds rich models of the world called cognitive maps that account for the relationships between locations, events, and experiences (e.g., Behrens et al., 2018; Moser, Kropff, & Moser, 2008; O’Keefe & Nadel, 1978; Tolman, 1948). Abstracting and organizing relational information in this way facilitates flexible behavior, enabling generalization and inference. Beyond classical findings on the importance of cognitive maps for spatial navigation (e.g., Burgess, Maguire, & O’Keefe, 2002; Ekstrom & Ranganath, 2018; O’Keefe & Nadel, 1978), they are also thought to organize the relationships between objects (Constantinescu, O’Reilly, & Behrens, 2016; Garvert, Dolan, & Behrens, 2017; Garvert, Saanum, Schulz, Schuck, & Doeller, 2021; Morton, Schlichting, & Preston, 2020, Theves, Fernandez, & Doeller, 2019, 2020; Viganò, Rubino, Di Soccio, Buiatti, & Piazza, 2021), to represent temporal distances (Bellmund, Deuker, & Doeller, 2019; Bellmund, Deuker, Montijn, & Doeller, 2022; Burgess, Maguire, & O’Keefe, 2002; Schapiro, Kustner, & Turk-Browne, 2012; Solomon, Lega, Sperling, & Kahana, 2019), and to structure knowledge in the context of social cognition (Park, Miller, Nili, Ranganath, & Boorman, 2020; Son, Bhandari, & Feldmanhall, 2021; Tavares et al., 2015). While cognitive mapping is thus proposed to be a universal, domain-unspecific coding principle to systematically organize knowledge (Behrens et al., 2018; Bellmund, Gärdenfors, Moser, & Doeller, 2018; Stachenfeld, Botvinick, & Gershman, 2017), it is unclear how the brain handles stimuli embedded in multiple relational structures that are very distinct in terms of their mode and timescale of acquisition. Does the hippocampal-entorhinal system form one conjunctive map that integrates similarities along the different stimulus dimensions or does it form anatomically separable maps for each stimulus dimension?

In Garvert et al. (2017), participants acquired new relational knowledge about everyday objects that were already embedded in semantic structures. Here, participants were trained on object sequences generated from a pseudo-random walk along a hidden graph. Within the hippocampal–entorhinal system, the neural representations of the presented objects reflected the link distance on the graph (Garvert et al., 2017). In this situation, besides the newly learned transitional probabilities between objects, participants can be assumed to have explicit knowledge about the semantic relationships between the same objects (e.g., rabbit and dog are both animals) as acquired over the course of their lifetime. We conjectured that prior semantic knowledge about objects would be simultaneously mapped in the same system, which also represents knowledge about transition probabilities.

Here we used a triplet odd-one-out task (Hebart, Zheng, Pereira, & Baker, 2020) to construct a model of object similarity. We matched the stimuli used in Garvert et al. (2017) with photographs of the same objects which have different visual features. This allowed us to isolate the semantic relationships, which reflect high-level conceptual knowledge acquired from experience, from low-level perceptual attributes of the specific objects (Rosch & Lloyd, 1978; Tversky, 1977), while maintaining the richness of the visually perceived stimuli. Using fMRI adaptation analysis on the data by Garvert et al. (2017), we found evidence consistent with a map of semantic relationships between objects that precisely localized to the hippocampus. Although both maps were represented in the hippocampal-entorhinal system, this semantic map was anatomically distinct from the previously described entorhinal map, which coded for newly learnt statistical regularities. By showing the hippocampal formation maps distinctive types of relationships simultaneously in parallel maps, our results thus demonstrate that the hippocampal formation does not form conjunctive maps that integrate similarities across distinct stimulus dimensions. Instead – at least in situations where the mode and timescale of acquisition are very distinct – different stimulus dimensions are organized in anatomically separable maps.

## Results

23 human participants were trained on object sequences whose transition probabilities followed a discrete, non-spatial graph (Figure 1A, Garvert et al. 2017). Without being consciously aware of the statistical regularities, participants’ neural activity in the hippocampus and entorhinal cortex reflected the transitional relationships they had experienced between the objects on a subsequent day (Figure 1B, 3A). However, the brain may not only represent the newly learned statistical regularities, but also the semantic relationships between the objects acquired over the course of a lifetime. Thus, we asked whether this information is also mapped in the same system.

**Figure 1.**
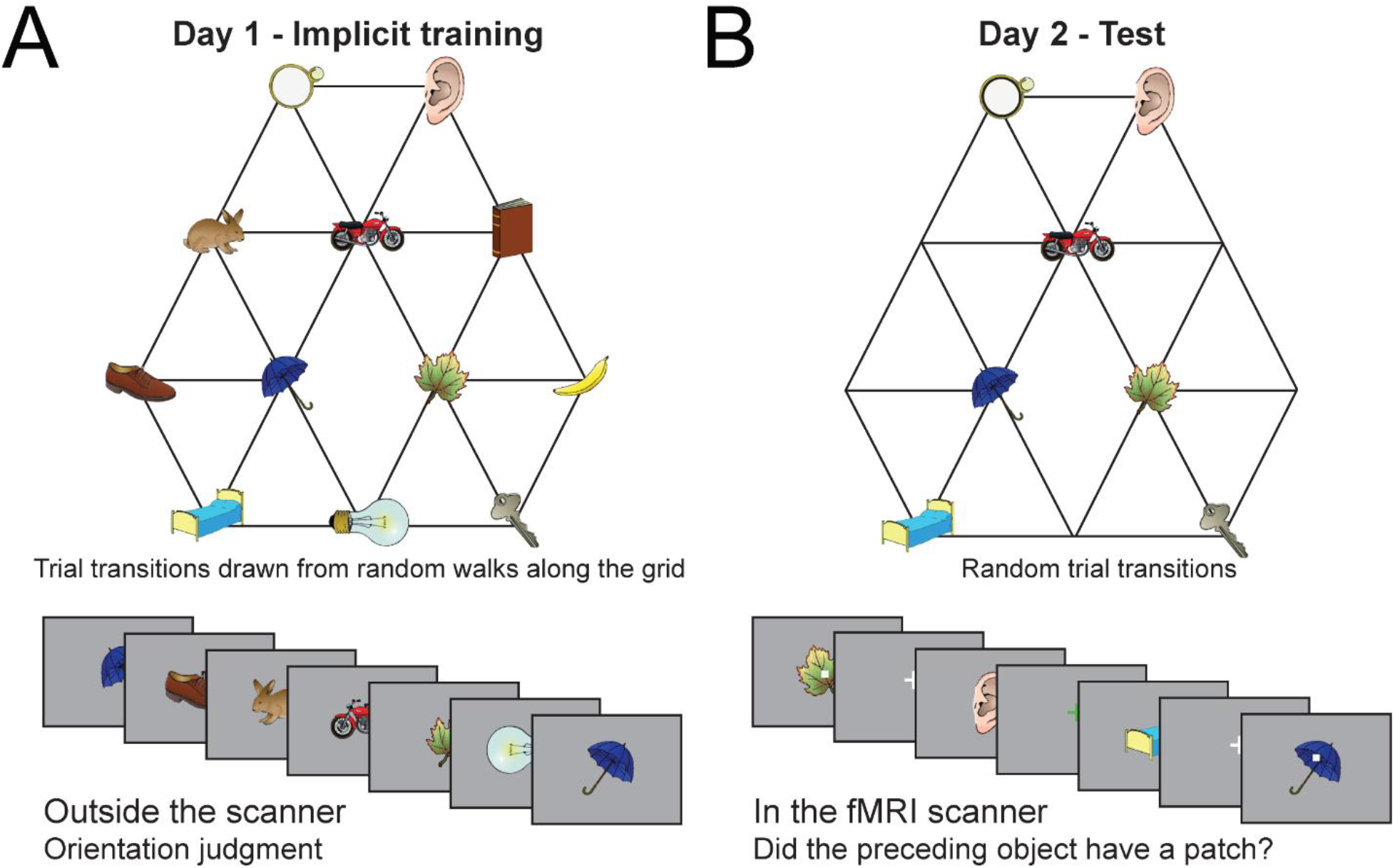
Experimental design. (A) Graph structure used to generate stimulus sequences on day 1. Trial transitions were drawn from random walks along the graph. (B) Objects on reduced graph presented to participants in the scanner on day 2. Trial transitions were random. In both sessions, participants performed simple behavioral cover tasks. Figure adapted from Garvert et al. (2017).

To address this question, we measured the semantic relationship between these objects using a triplet odd-one out task, where participants were shown three objects on each trial and asked to select the image that was the least similar to the other two (Figure 2A, Hebart et al., 2020). By repeatedly varying the third object for a pair of target objects, their similarity could be assessed in a wide range of different contexts. We first asked a separate group of 128 participants online to rate the similarities of all 31 objects used in the original study. We then computed object similarity as the probability of participants choosing two objects together, irrespective of the context imposed by the third object. To separate the semantic relationships between objects from the perceptual similarities, we matched our objects (Figure 2B, top rows) to corresponding real-world photographs of the same objects in the THINGS database (Figure 2B, bottom rows; Hebart et al., 2019). The THINGS database shows photographs of objects embedded in a natural environment as opposed to simple line drawings of stereotypical objects used in our fMRI study. In this way, the shared perceptual similarity between objects in these two data sets should be reduced. The similarity of objects in the THINGS database was assessed by 5,301 participants in an independent study using the same triplet odd-one-out task, in the context of a total of 1,854 objects (Hebart et al., 2020). The similarity ratings obtained for our objects and the corresponding objects in the THINGS database were highly correlated (Spearman’s Rho = .89, *p* < .001, Figure 2C) as expected if semantic relations are preserved across datasets.

**Figure 2.**
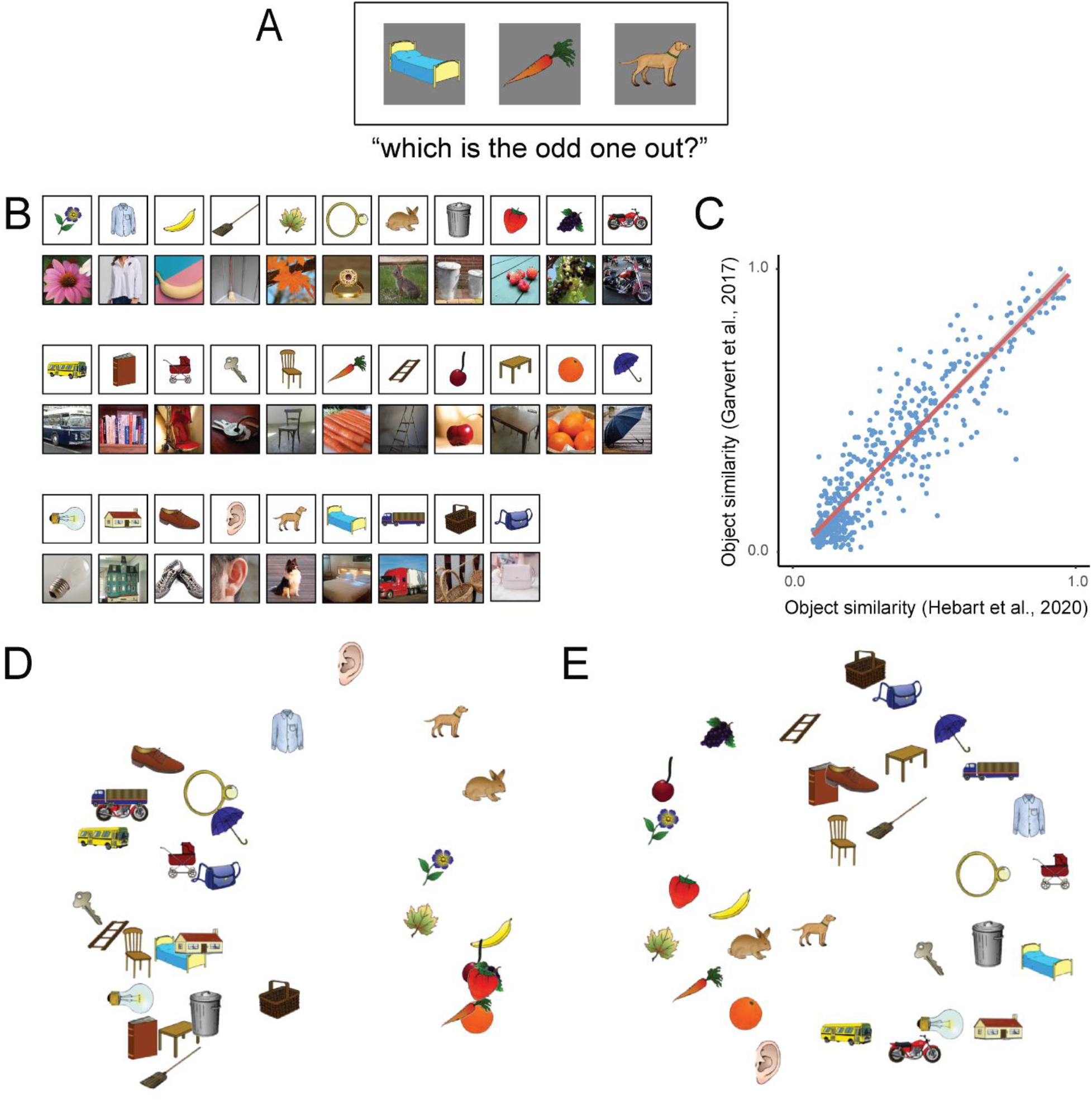
Semantic distance constructed using the triplet odd-one-out task. (A) An example trial of the triplet odd-one-out task. The task measures object similarity as the probability of participants choosing two objects together, irrespective of the context imposed by the third object (Hebart et al., 2020). (B) Stimuli used in the odd-one-out task. Top rows: all 31 stimuli from the original study; bottom rows: a subset of stimuli from the THINGS database, matched with the 31 object stimuli used in the original study. The rating of the matched objects is done in the context of a total of 1,854 objects (Hebart et al., 2020). (C) Correlation between similarity ratings based on our own stimuli and ratings based on the corresponding stimuli from the THINGS database (Spearman’s Rho = .70, *p* < .001). (D) Visualization of the 31 objects’ semantic distance in a two-dimensional space according to multi-dimensional scaling (MDS). (E) 2D MDS visualization of the 31 objects’ residual distance.

We regressed the matched similarity matrix (x-axis, Figure 2C) onto our original similarity matrix (y-axis, Figure 2C). By doing this, we were able to separate the variance into two parts: (1) the part that could be explained by an independent measure of object similarity obtained from a matched set of stimuli, and (2) the part that could not be explained by the independent measure. Although semantic and perceptual features are unlikely to be fully separable in this way, we consider the first part to primarily reflect the semantic relationships between our objects that are preserved across different ways of visualizing objects (hereafter: semantic distance); while the second part reflected a combination of features that are not accounted for in terms of semantics, including perceptual similarities (hereafter: residual distance). To visualize the most faithful two-dimensional representation of distances between objects, we applied multidimensional scaling (MDS) to the two resulting distance matrices. The semantic MDS reveals that the similarity ratings led to the emergence of object category clusters (e.g., fruit, animals, man-made objects) and replicates well-known distinctions between “animate-inanimate” and “natural - man-made” (Hebart et al. 2020, Figure 2D). The residual distance continued to express differences between man-made and natural objects, however the overall arrangement was less structured (Figure 2E). Neither the semantic distance nor the residual distance was correlated with link distance (semantic: Spearman’s Rho mean = .03, SD = .12, range = -.25 – .30, *t*_22_ = 1.04, *p* = .31); residual: Spearman’s Rho mean = -.03, SD = .09, range = -.22 – .15, *t*_22_ = -1.52, *p* = .14).

Following the approach adopted in the original study, we exploited fMRI adaptation (Barron, Garvert, & Behrens, 2016; Grill-Spector, Henson, & Martin, 2006) to investigate the representational similarity for different objects. fMRI adaptation relies on the observation that the repeated activation of the same population of neurons leads to a suppressed response. In this way, the amount of suppression can serve as a proxy for the similarity of the underlying neural representations. In line with the decrease in fMRI adaptation as a function of link distance observed in the entorhinal cortex (Garvert et al. 2017), we reasoned that in areas representing object relationships (e.g., semantic relationships), fMRI adaptation should scale with the corresponding distance measures (i.e., semantic distance). We included link distances, semantic distance, and residual distance as parametric regressors in the same GLM and looked for brain regions whose fMRI responses to each object decreased as a linear function of these distance measures to the preceding object.

We expected both the semantic information and the statistical regularities to be mapped in the hippocampal formation. Therefore, we focused our analysis on regions within an anatomical mask comprising bilateral entorhinal cortex, subiculum, and hippocampus (see mask used for small-volume correction in Supplementary material S1). Areas outside these regions were only considered significant if they survived whole-brain correction.

The results replicated our original finding of the link distance effect after accounting for the semantic distance and the residual distance. The fMRI adaptation analysis showed a cluster bilaterally in the entorhinal cortex (Figure 3A), with the left, but not the right peak surviving small-volume correction within the entorhinal-hippocampal mask (family-wise error (FWE) corrected at peak level, left peak *t*_22_ = 4.44, *p* = .029, [-18, -19, -25], right peak *t*_22_ = 3.50, *p* = .17, [21, -22, - 25]). Critically, we also observed a semantic distance effect in the bilateral hippocampus (Figure 3B), which was significant in the right hemisphere in the same entorhinal-hippocampal mask (left peak *t*_22_ = 4.10, *p* = .06, [-27, -34, -10], right peak *t*_22_ = 4.69, *p* = .019, [24, -31, -10]). No region showed fMRI adaptation effects covarying with residual distance (all *p*s > .99, FWE corrected on the cluster level, no suprathreshold cluster).

**Figure 3.**
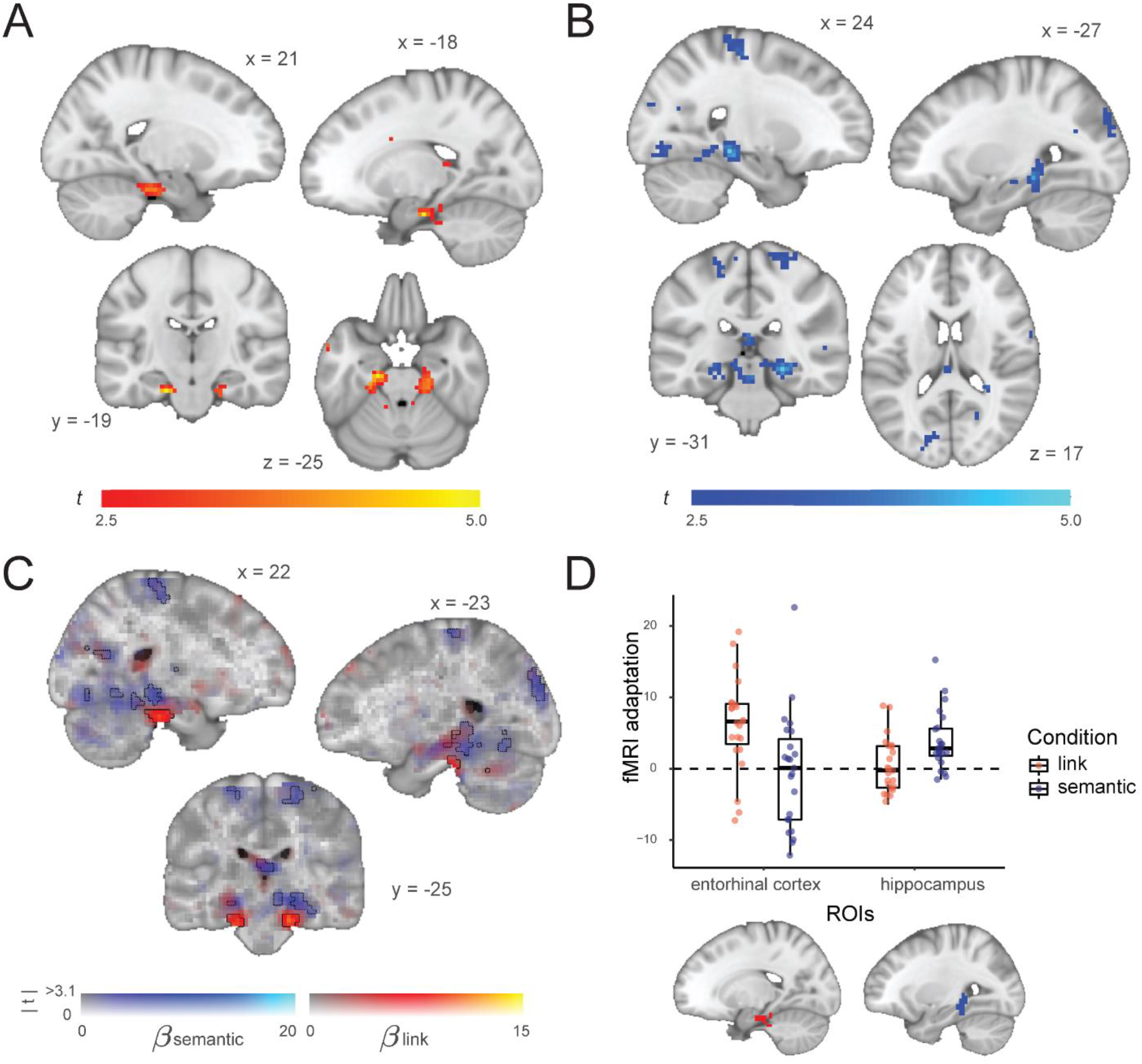
Statistical and semantic relationships are represented in non-overlapping clusters in the hippocampal-entorhinal system. (A) Whole-brain analysis showing a decrease in fMRI adaptation with link distance in the entorhinal cortex, when link distance, semantic distance and residual distance are included in the model. (B) Whole-brain analysis showing a decrease in fMRI adaptation with semantic distance in the hippocampus, when link distance, semantic distance and residual distance are included in the model. (C) Link distance effect (red) and semantic distance effect (blue) are represented in non-overlapping clusters. Whole-brain results are displayed using Slice Display (Zandbelt, 2017) using a dual-coding data visualization approach (Allen, Erhardt, & Calhoun, 2012), with color indicating the parameter estimates and opacity the associated *t* statistics. Solid and dotted contours outline statistically significant clusters for the link and the semantic effects, respectively. (D) Bottom: The two ROIs defined based on the link distance effect in the entorhinal cortex (in red) and the semantic distance effect in the hippocampus (in blue). Top: boxplot of the parameter estimates for the link distance and semantic distance effects extracted from the two ROIs. The thick horizontal line inside the box indicates the median, and the bottom and top of the box indicate the first and third quartiles of each condition. Each dot represents one participant. The plot is for visualization only, since the contrast used for defining the ROIs is not independent from the interaction effect of interest here. Both (A) and (B) are thresholded at *p* < .01, uncorrected for visualization.

When we assessed object dissimilarity using our own stimuli, a measure that encompasses not only semantic, but also perceptual and other types of object relationships, and used this as a predictor in our fMRI adaptation analysis, we identified a cluster in the visual cortex, whereas a hippocampal cluster surpassed the cluster-defining threshold, but did not survive correction for multiple comparisons (Supplementary Material S2). This is consistent with the view that object similarity ratings are strongly influenced by both perceptual and semantic features (Rosch & Lloyd, 1978; Tversky, 1977).

While the link distance and the semantic distance are both represented in the hippocampal formation, they were located in two non-overlapping clusters within this region (Figure 3C). To investigate this at a more fine-grained level, we defined two functional regions of interest (ROIs): (1) a bilateral ROI in the entorhinal cortex defined by the link distance effect (Figure 3A, hereafter: EC ROI), and (2) a bilateral ROI in the hippocampus defined by the semantic distance effect (Figure 3B, hereafter: HC ROI). We used these two ROIs (both thresholded at *p* < .01, uncorrected) to extract parametric estimates from the opposite contrast. As shown in Figure 3D, the individual semantic distance effect extracted from the EC ROI and the link distance effect extracted from the HC ROI were not significantly different from 0 (semantic: *t*_22_ = -0.25, *p* = .81; link: *t*_22_ = 0.54, *p* = .60).

Together, our results suggest that both recently learned statistical regularities of which participants have no conscious awareness, as well as long-term semantic relationships that are explicitly accessible and acquired over the course of a lifetime, are represented in the hippocampal formation simultaneously, albeit in different subregions: While transition probabilities are represented in more entorhinal regions, semantic relationships are represented in more hippocampal regions.

## Discussion

The brain forms cognitive maps of the relationships between landmarks that help an animal navigate their physical environment (Burgess et al., 2002; Ekstrom & Ranganath, 2018; O’Keefe & Nadel, 1978; Tolman, 1948). Previous studies have shown that the same organizing principle also applies to other non-spatial types of relational information (Constantinescu et al., 2016; Garvert et al., 2017; Garvert et al. 2021; Morton et al., 2020; Theves et al, 2019, 2020; Viganò et al, 2021). For example, when participants acquire new knowledge about the relationships between objects by being exposed to experimentally generated object sequences, the hippocampal formation extracts the associated transition structure and stores it as a cognitive map (Garvert et al. 2017). However, it is also the case that participants already have existing knowledge about the semantic relationships between these objects acquired over an entire lifetime. Here we show that this prior knowledge is also simultaneously mapped in the same neural system that codes for newly learned structural information. This suggests a common framework for the representation of relational knowledge that extends beyond relationships that can be extracted from the statistics of sequences we experience in the short term, with knowledge acquired over vastly different time frames (an hour on the previous day as opposed to an entire lifetime) organized in similar ways.

Cognitive mapping is proposed to be an organizing principle that underlies our ability to generalize and make inferences (Behrens et al., 2018). Here we show that the hippocampal formation extracts the embedding of a stimulus in multiple relational structures even when neither stimulus feature is directly task relevant. The representation of both a statistical map and a semantic map in the same system is remarkable, given their very different timescales and modes of acquisition (i.e., implicit vs. explicit learning). However, while the two relational structures are represented in the same neural system, they are not represented in overlapping voxels, suggesting that the brain extracts separable relational structures in parallel rather than integrating multiple structures in one compositional map (Spiers, 2020). A representation of separable maps could be useful in situations where different stimulus dimensions can become relevant for generalization at different times, enabling the hippocampus to guide generalization flexibly depending on the task demands (Garvert et al., 2021). Our finding that a semantic map is represented in the hippocampus is also consistent with previous findings that hippocampal activity reflects distances in semantic spaces (Estefan et al., 2021; Romero, Barense, & Moscovitch, 2019; Solomon et al., 2019). Notably, we decoded the semantic map even though we did not explicitly manipulate the semantic similarity of the stimuli and this information was not task-relevant, demonstrating how prevalent the representation of relational information is in the hippocampus.

Interestingly, the statistical map is found in the entorhinal cortex whereas the semantic map is represented in the hippocampus. One potential reason for this segregation may relate to a difference in the “age” of the acquired regularities. Whereas the statistical regularities were learnt on the day prior to scanning, the semantic relationships were acquired over the course of one’s lifetime. One possibility is that relational memories might transition from the entorhinal cortex (“new” knowledge) to the hippocampus (“old” knowledge), a suggestion that however contradicts the conventional view that memories are first encoded in hippocampus and later represented in the neocortex in a more permanent form of storage (Dudai, Karni, & Born, 2015; Marr, 1971). Alternatively, the segregation of the two maps might reflect differences in the learning process by which statistical versus semantic relational knowledge is acquired. The formation of a statistical map requires learning transitional probabilities between sequential states (Whittington, McCaffary, Bakermans, & Behrens, 2022). Entorhinal grid cell activity has been suggested to reflect a principal component decomposition of predictive maps, or successor representations (Stachenfeld, Botvinick, & Gershman, 2014; Stachenfeld et al., 2017). A representation of the transition structure in this region is consistent with this account of grid cell function. Conversely, semantic knowledge is not necessarily acquired in a sequential fashion and might instead reflect common associations or co-occurrences between objects, a representation much better suited to putative coding principles underlying hippocampal place cell firing.

In sum, our study shows that the hippocampal-entorhinal system extracts diverse relational structures in which a stimulus is embedded. Both the semantic and the statistical maps are separately and simultaneously represented even when neither structure is task relevant. This allows relevant knowledge to be flexibly selected at a later timepoint in order to guide goal-directed behavior in novel situations (Behrens et al., 2018; Spiers, 2020; Whittington et al., 2020).

## Materials and Methods

### fMRI study

23 human participants (15 men, *mean*_age_ = 23.5, *SD*_age_ = 3.7, *range*_age_ 18-31) were trained on object sequences whose transitions followed a pseudo-random walk along a graph (Figure 1A). The graph structure was the same for all participants, with the link distance between objects on the graph ranging from one to four. For each participant, a subset of 12 objects were selected from a total of 31 objects used in the study, and randomly assigned to the 12 nodes on the graph. Objects within the same semantic categories were avoided to be assigned to the same participant. In the scanning session, 7 out of the 12 training objects were used (Figure 1B) to reduce the total number of stimulus–stimulus transitions and thereby increase statistical power.

### Behavioral experiment of object similarity

#### Participants

A separate group of 128 workers from the online crowdsourcing platform Amazon Mechanical Turk took part in a triplet odd-one-out task (55 men, *mean*_age_ = 42.6, *SD*_age_ = 11.9, *range*_age_ 20-70). All workers were located in the United States and provided informed consent. The online research was approved by the Office of Human Research Subject Protection and conducted following all relevant ethical regulations, and the workers were compensated financially for their time.

#### Stimuli and task

The 31 objects in the original study (Garvert et al., 2017) were used in the triplet odd-one-out task. All images depict colored and shaded objects and were selected from the “Snodgrass and Vanderwart” database (Rossion & Pourtois, 2004).

The task was carried out in sets of 20 trials. Participants could choose how many sets they would like to take part in. On average, participants performed 140.63 trials (*min* = 20, *max* = 1460), with a median RT of 2221 ms. On each trial, participants were shown three object images side by side and were asked to select the image that was the least similar to the other two (Figure 2A). Each object triplet and the order of stimuli were chosen randomly, but such that after collection of the entire data set each cell in the 31 × 31 similarity matrix had been sampled at least once. The object similarity was defined as the probability *p*(i,j) of the participants choosing objects i and j to belong together, irrespective of context (Hebart et al., 2020).

### fMRI data analysis

#### Computation of the parametric regressors

For each participant, we compute a link distance matrix and three matrices describing object relations (i.e., object dissimilarity, semantic distance, residual distance, explained below). Whereas the link distance matrix (values range from 1 to 4) was identical for all participants (Figure S2A, left panel), the object similarity matrices were unique for each participant (Figure S2A, right panel). To turn object similarities into a distance measure, we computed object dissimilarities by subtracting the similarity values from 1. Therefore, a number close to 1 means that two objects are dissimilar to each other, whereas a number close to 0 means the objects are very similar to each other.

The object dissimilarity matrix was derived directly from the triplet odd-one-out task described above. Given that each participant received a different set of object stimuli in the training, we standardized the individual 12 * 12 dissimilarity matrices for each participant and used the *z*-score values as parametric regressors in the fMRI analysis.

To isolate semantic relationships, we made use of an independent similarity rating from a different dataset (Hebart et al., 2020). The rationale is that the part of variance that can be explained by an independent rating reflects prior semantic knowledge that is independent of the precise visual display of a particular object. The independent rating is based on 1,854 images from the THINGS database (Hebart et al., 2019) which depicts photographs of objects embedded in a natural background, rated by a total of 5,301 participants using the same triplet odd-one-out task. From the 1,854 images, we selected 31 pictures (Figure 2B) that depict the same objects as the 31 objects in the original study of Garvert et al. (2017) and computed a sub-matrix of these 31 objects. Since each fMRI participant was trained on 12 out of the total 31 stimuli, we linearly regressed for each participant the 12 × 12 dissimilarity matrix based on object images from the THING database (X in the regression below, Hebart et al., 2020) onto the 12 × 12 dissimilarity matrix based on object images from the original fMRI experiment (Y in the regression, Garvert et al., 2017). Both matrices were *z*-scored, therefore no intercept was included in the regression.

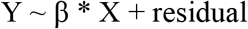

We consider β * X to reflect the variance of object dissimilarity that could be explained by an independent rating, likely to capture mostly semantic relationships. In contrast, the residual values reflect the variance that is not shared across different visual object stimuli, including specific perceptual features.

To visualize the object relatedness acquired from the triplet odd-one-out task, we performed multidimensional scaling (MDS) on the dissimilarity matrices. In the output MDS (Figures 2D, 2E, S2B), objects are arranged on a two-dimensional space, where the Euclidean distances reflect the dissimilarities between objects as well as possible. Note that MDS can only be performed on matrices with positive entries. We therefore subtracted the minimum value of the matrix and added 1. The addition of constants does not affect the resulting visualization of distances.

#### fMRI adaptation analysis

We performed two event-related generalized linear models (GLMs) to analyze the fMRI data. In the GLM for the main analysis, we included three parametric regressors which corresponded to the link distance (defined as the minimum number of links between the pairs of objects; i.e. distance 1, 2, or 3) and the semantic and residual distance derived from the ratings of the triplet odd-one-out task between the object on trial t and the preceding object on trial t-1. In the GLM for the supplementary analysis, we included the object dissimilarity regressor extracted from the odd-one-out task performed on the original stimuli and the link distance regressor. All regressors were standardized in the GLMs. No orthogonalization was applied.

Both GLMs contained separate onset regressors for each of the seven objects with a patch and without a patch. Each onset regressor was accompanied by different parametric regressors. By analogy to the original analysis, both GLMs included a button press regressor as a regressor of no interest. Trials associated with a button press and the two subsequent trials were not included in the main regressors in order to avoid button press-related artifacts. The same six motion regressors and the 17 physiological regressors (ten for cardiac phase, six for respiratory phase and one for respiratory) used in the original analysis were also included in the current GLMs. All regressors were convolved with a canonical hemodynamic response function. Blocks were modeled separately within the GLMs. Only the non-patch trials were included for our contrasts of interest. The contrast images of all participants from the first level were analyzed as a second-level random effects analysis. We report our results at a cluster-defining statistical threshold of *p* < .01 uncorrected, combined with small-volume correction (SVC) for multiple comparisons (peak-level FWE corrected at *p* < .05). For the SVC procedure, we used an anatomically defined mask comprising bilateral entorhinal cortex, subiculum, and hippocampus (Supplementary figure S1). Activations in other brain regions were only considered significant at a level of *p* < .001 uncorrected if they survived whole-brain FWE correction at the cluster level (*p* < .05). All statistical parametric maps presented in the manuscript are unmasked.

To illustrate the non-overlapping clusters for the link distance effect and the semantic distance effect (Figure 3C), we defined two regions of interest (ROIs) based on the two parametric estimations from non-patch trials in the main GLM. The link distance effect revealed a cluster in bilateral entorhinal cortex, which we used to define the EC ROI; the semantic distance effect revealed a cluster in bilateral hippocampus, which we used to define the HC ROI. For both ROIs we included all the voxels exceeding a *t* value of 2.5, corresponding to *p* < .01. From the two ROIs, we then extracted parameter estimates for each of the 23 participants for the two effects (Figure 3D). Due to the statistical dependence between the data and the ROI definition, no statistical inference was made regarding the interaction.

The statistical analysis was done using SPM12 (Wellcome Trust Centre for Neuroimaging, http://www.fil.ion.ucl.ac.uk/spm) and visualization was done using FSLeyes (Wellcome Centre for Integrative Neuroimaging, https://git.fmrib.ox.ac.uk/fsl/fsleyes/fsleyes/). The dual coded visualization of the fMRI data (Figure 3C) used a procedure introduced by Allen et al. (2012) and implemented by Zandbelt (2017). Clusters in the dual-coded map show brain activities as a function of the link distance (red) and the semantic distance (blue) simultaneously. The hue indexes the size of the parameter estimate, and the opacity indexes the unthresholded *t* values. Significant clusters (cluster-level corrected, FWE, *p* < .05) are encircled in solid (link distance) or dotted (semantic distance) contours.

## Data and Code Availability

Data and codes of the original study are available on datadryad (https://doi.org/10.5061/dryad.nk08s).

Additional data and codes for the current follow-up study are available at the Donders Repository (https://doi.org/10.34973/8m6q-qj39). They will be shared publicly upon manuscript acceptance.

## Acknowledgements

This study was supported by the Dutch Research Council (NWO) under Gravitation grant number (024.001.006) to the Language in Interaction Consortium.

## Supplementary Materials

**Supplementary Figure S1:**
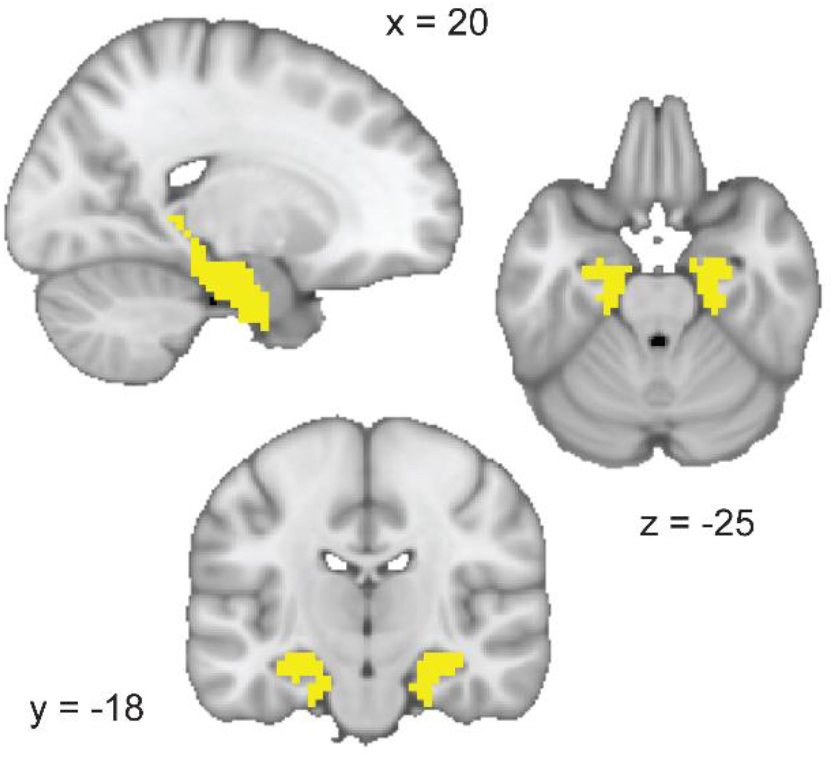
Anatomically defined mask used for small-volume correction (incl. bilateral entorhinal cortex, subiculum, and hippocampus)

### Supplementary Material S2: Additional analysis of object similarity

To explore the representation of stimulus dissimilarities for the objects we used in our study, we ran a triplet odd-one-out task (Hebart et al., 2020) in a separate group of 128 participants. Participants were instructed to select the object that was the least similar to the other two, without a specific instruction as to whether specific stimulus features such as semantic or perceptual similarities should guide this decision. Stimulus similarity ratings are therefore likely to be influenced by both of those stimulus dimensions. We then computed object similarity as the probability of participants choosing two objects together, irrespective of the context imposed by the third object and transformed the similarity rating into an object dissimilarity measure by subtracting each similarity measure from 1. Since every participant was only exposed to a unique subset consisting of 12 of the 31 objects, the dissimilarity matrix was different for each participant (Figure S2A, right panel). In contrast, the link distance between objects on the graph was the same for all participants (Figure S2A, left panel). It is worth noting that the link distance matrix and the object dissimilarity matrix did not correlate across participants (Spearman’s Rho *mean* = .01, *SD* = .11, *range* = -.20 – .25, *t*_22_ = 0.34, *p* = .74). We standardized each individual’s 12 * 12 object dissimilarity matrix based on the unique set of 12 objects each participant was trained on. A two-dimensional representation of distances between objects in the dissimilarity matrix using MDS confirms that the objects are closely grouped according to semantic features (e.g., natural - man-made, Figure S2B). In addition, perceptual properties such as object color also influenced similarity ratings to some degree, suggesting that object similarity ratings reflect a mixture of perceptual and semantic relationships (Rosch & Lloyd, 1978; Tversky, 1977).

To investigate the representation of these dissimilarity structures in the brain, we set up a GLM that included the link distance as well as this measure of object dissimilarity. We observed that fMRI adaptation scaled as a function of object dissimilarity in the visual cortex (FWE corrected at cluster-level, K_E_ = 86, *p* = .004, peak *t*_22_ = 5.49, [27, -76, 17]). Voxels in the hippocampal formation also passed the cluster-defining threshold for both the object dissimilarity effect (Figure S2C) and the link distance effect (Figure S2D). However, neither of them survived small-volume correction using the hippocampal-entorhinal mask (link: right peak *t*_22_ = 3.93, *p* = .08, [24, -25, -25] and left peak *t*_22_ = 3.40, *p* = .21, [-18, -19, -25]; object: right peak *t*_22_ = 4.01, *p* = .07, [24, -31, -10] and left peak *t*_22_ = 3.42, *p* = .21, [-24, -31, -10]). We reason that this might be due to similarity ratings in this setting reflecting a mixture of semantic and perceptual similarities.

**Supplementary Figure S2.**
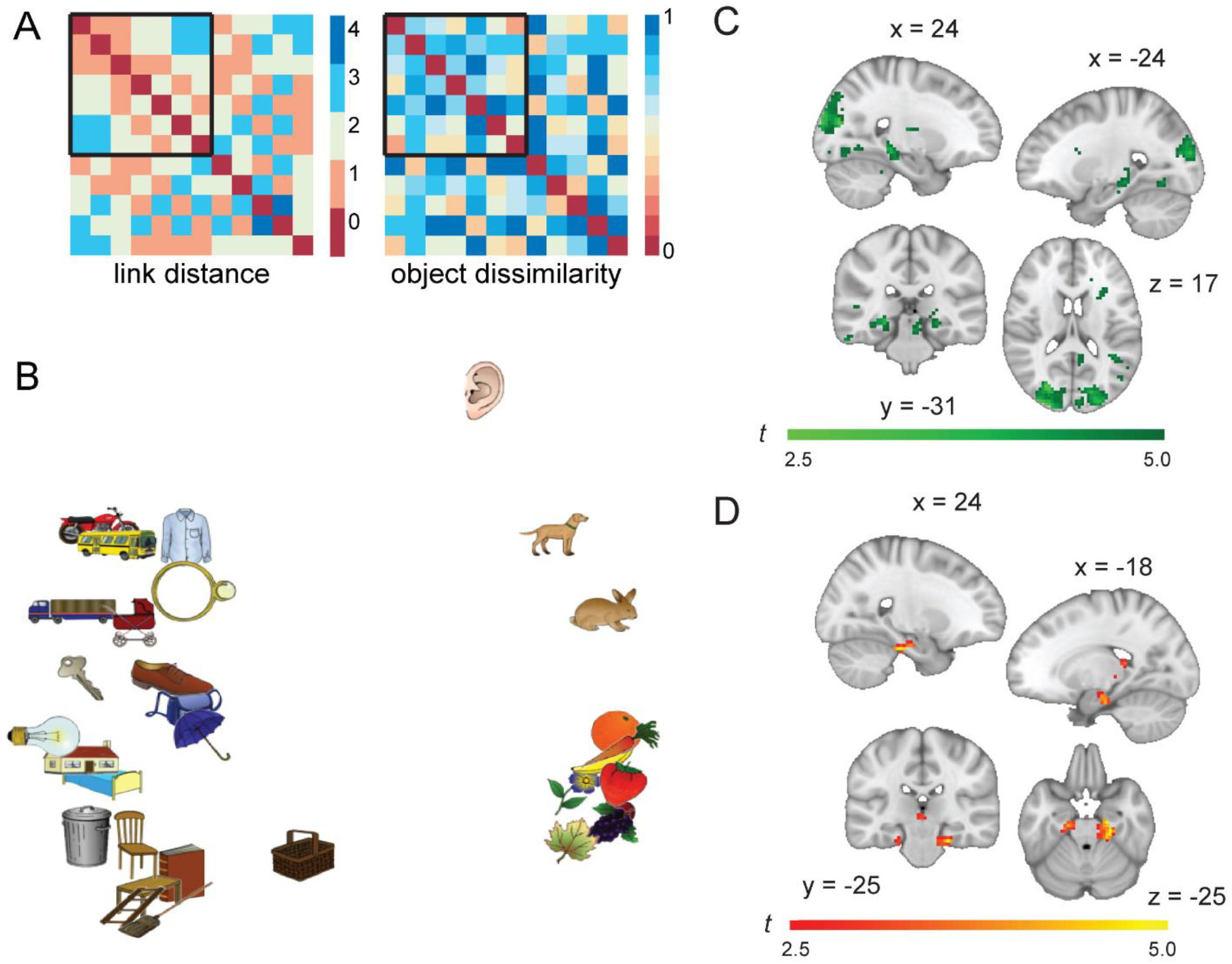
Additional analysis of object similarity. (A) The link distance matrix (left) and one example object dissimilarity matrix (right) corresponding to the graph structure and the stimulus set used in the original fMRI study. The link distance matrix was the same for all participants. The object dissimilarity matrix was unique for each participant. The black frame indicates the subset of seven objects presented in the scanner and used in the fMRI analysis. (B) Visualization of the object dissimilarity matrix in two dimensions, according to multi-dimensional scaling (MDS). More similar objects are located near each other. (C, D) fMRI adaptation effects of object dissimilarity (C) and link distance (D). Whole-brain analysis showing a decrease in fMRI adaptation with object dissimilarity in the visual cortex, when link distance and object dissimilarity are both included as parametric modulators in the model. All images are thresholded at *p* < .01, uncorrected for visualization.

## References

Allen, E. A., Erhardt, E. B., & Calhoun, V. D. (2012). Data visualization in the neurosciences : Overcoming the curse of dimensionality. Neuron, 74(4), 603–608. https://doi.org/10.1016/j.neuron.2012.05.001

Barron, H. C., Garvert, M. M., & Behrens, T. E. J. (2016). Repetition suppression: A means to index neural representations using BOLD? Philosophical Transactions of the Royal Society B: Biological Sciences, 371(1705). https://doi.org/10.1098/rstb.2015.0355

Behrens, T. E. J., Muller, T. H., Whittington, J. C. R., Mark, S., Baram, A. B., Stachenfeld, K. L., & Kurth-Nelson, Z. (2018). What is a cognitive map? Organizing knowledge for flexible behavior. Neuron, 100, 490–509. https://doi.org/10.1016/j.neuron.2018.10.002

Bellmund, J. L. S., Gärdenfors, P., Moser, E. I., & Doeller, C. F. (2018). Navigating cognition: Spatial codes for human thinking. Science, 362. https://doi.org/10.1126/science.aat6766

Bellmund, J. L., Deuker, L., & Doeller, C. F. (2019). Mapping sequence structure in the human lateral entorhinal cortex. eLife, 8, e45333. https://doi.org/10.7554/eLife.45333

Bellmund, J. L., Deuker, L., Montijn, N. D., & Doeller, C. F. (2022). Mnemonic construction and representation of temporal structure in the hippocampal formation. Nature Communications, 13(10), 1–16. https://doi.org/10.1038/s41467-022-30984-3

Burgess, N., Maguire, E. A., & O’Keefe, J. (2002). The human hippocampus and spatial and episodic memory. Neuron, 35(4), 625–641. https://doi.org/10.1016/S0896-6273(02)00830-9

Constantinescu, A. O., O’Reilly, J. X., & Behrens, T. E. J. (2016). Organizing conceptual knowledge in humans with a gridlike code. Science, 352, 1464–1468. https://doi.org/10.1126/science.aaf0941

Dudai, Y., Karni, A., & Born, J. (2015). The consolidation and transformation of memory. Neuron, 88, 20–32. https://doi.org/10.1016/j.neuron.2015.09.004

Ekstrom, A. D., & Ranganath, C. (2018). Space, time, and episodic memory: The hippocampus is all over the cognitive map. Hippocampus, 28(9), 680–687. https://doi.org/10.1002/hipo.22750

Estefan, D. P., Zucca, R., Arsiwalla, X., Principe, A., Zhang, H., Rocamora, R., Axmacher, N., & Verschure, P. F. M. J. (2021). Volitional learning promotes theta phase coding in the human hippocampus. Proceedings of the National Academy of Sciences of the United States of America, 118(10), 1–12. https://doi.org/10.1073/pnas.2021238118

Garvert, M. M., Dolan, R. J., & Behrens, T. E. J. (2017). A map of abstract relational knowledge in the human hippocampal–entorhinal cortex. ELife, 6, 1–20. https://doi.org/10.7554/eLife.17086

Garvert, M. M., Saanum, T., Schulz, E., Schuck, N. W., & Doeller, C. F. (2021). Hippocampal spatio-temporal cognitive maps adaptively guide reward generalization. bioRxiv. https://doi.org/10.1101/2021.10.22.465012

Grill-spector, K., Henson, R., & Martin, A. (2006). Repetition and the brain : neural models of stimulus-specific effects. Trends in Cognitive Science, 10(1), 17–19. https://doi.org/10.1016/j.tics.2005.11.006

Hebart, M. N., Dickter, A. H., Kidder, A., Kwok, W. Y., Corriveau, A., Wicklin C. Van, & Baker, C. I. (2019). THINGS : A database of 1,854 object concepts and more than 26,000 naturalistic object images. PLoS One, 14(10), e0223792. https://doi.org/https://doi.org/10.1371/journal.pone.0223792

Hebart, M. N., Zheng, C. Y., Pereira, F., & Baker, C. I. (2020). Revealing the multidimensional mental representations of natural objects underlying human similarity judgements. Nature Human Behaviour, 4(11), 1173–1185. https://doi.org/10.1038/s41562-020-00951-3

Marr, D. (1971). Simple memory: a theory for archicortex. Philosophical Transactions of the Royal Society B, 262, 23–81. https://doi.org/10.1098/rstb.1971.0078.

Morton, N. W., Schlichting, M. L., & Preston, A. R. (2020). Representations of common event structure in medial temporal lobe and frontoparietal cortex support efficient inference. Proceedings of the National Academy of Sciences of the United States of America, 117(47), 29338–29345. https://doi.org/10.1073/pnas.1912338117

Moser, E. I., Kropff, E., & Moser, M. B. (2008). Place cells, grid cells, and the brain’s spatial representation system. Annual Review of Neuroscience, 31, 69–89. https://doi.org/10.1146/annurev.neuro.31.061307.090723

O’Keefe, J., & Nadel, L. (1978). The Hippocampus as a Cognitive Map. Clarendon Press. https://doi.org/10.5840/philstudies19802725

Park, S. A., Miller, D. S., Nili, H., Ranganath, C., & Boorman, E. D. (2020). Map Making: Constructing, Combining, and Inferring on Abstract Cognitive Maps. Neuron, 1–13. https://doi.org/10.1016/j.neuron.2020.06.030

Romero, K., Barense, M. D., & Moscovitch, M. (2019). Coherence and congruency mediate medial temporal and medial prefrontal activity during event construction. NeuroImage, 188(August 2018), 710–721. https://doi.org/10.1016/j.neuroimage.2018.12.047

Rosch, E., & Lloyd, B. B. (1978). Cognition and categorization. John Wiley & Sons.

Rossion, B., & Pourtois, G. (2004). Revisiting Snodgrass and Vanderwart ‘ s object pictorial set : The role of surface detail in basic-level object recognition À. Perception, 33, 217–237. https://doi.org/10.1068/p5117

Schapiro, A. C., Kustner, L. V., & Turk-Browne, N. B. (2012). Shaping of object representations in the human medial temporal lobe based on temporal regularities. Current Biology, 22(17), 1622–1627. https://doi.org/10.1016/j.cub.2012.06.056

Solomon, E. A., Lega, B. C., Sperling, M. R., & Kahana, M. J. (2019). Hippocampal theta codes for distances in semantic and temporal spaces. Proceedings of the National Academy of Sciences of the United States of America, 116(48), 24343–24352. https://doi.org/10.1073/pnas.1906729116

Son, J., Bhandari, A., & Feldmanhall, O. (2021). Cognitive maps of social features enable flexible inference in social networks. PNAS, 1–11. https://doi.org/10.1073/pnas.2021699118

Spiers, H. J. (2020). The Hippocampal Cognitive Map: One Space or Many? Trends in Cognitive Sciences, 24(3), 168–170. https://doi.org/10.1016/j.tics.2019.12.013

Stachenfeld, K. L., Botvinick, M. M., & Gershman, S. J. (2014). Design principles of the hippocampal cognitive map. In Advances in Neural Information Processing Systems, 27.

Stachenfeld, K. L., Botvinick, M. M., & Gershman, S. J. (2017). The hippocampus as a predictive map. Nature Neuroscience, 20, 1643–1653. https://doi.org/10.1038/nn.4650

Tavares, R. M., Mendelsohn, A., Grossman, Y., Williams, C. H., Shapiro, M., Trope, Y., & Schiller, D. (2015). A Map for Social Navigation in the Human Brain. Neuron, 87(1), 231–243. https://doi.org/10.1016/j.neuron.2015.06.011

Theves, S., Fernandez, G., & Doeller, C. F. (2019). The hippocampus encodes distances in multidimensional feature space. Current Biology, 29(7), 1226–1231. https://doi.org/10.1016/j.cub.2019.02.035

Theves, S., Fernández, G., & Doeller, C. F. (2020). The hippocampus maps concept space, not feature space. Journal of Neuroscience, 40(38), 7318–7325. https://doi.org/10.1523/JNEUROSCI.0494-20.2020

Tolman, E. C. (1948). Cognitive maps in rats and men. Psychological Review, 55, 189–208. https://doi.org/10.1037/h0061626

Tversky, A. (1977). Features of Similarity. Psychological Review, 84(4), 327–352. https://doi.org/10.1037/0033-295X.84.4.327

Viganò, S., Rubino, V., Di Soccio, A., Buiatti, M., & Piazza, M. (2021). Grid-like and distance codes for representing word meaning in the human brain. NeuroImage, 232, 117876. https://doi.org/10.1016/j.neuroimage.2021.117876

Whittington, J. C. R., Muller, T. H., Mark, S., Chen, G., Barry, C., Burgess, N., & Behrens, T. E. J. (2020). The Tolman-Eichenbaum Machine: Unifying Space and Relational Memory through Generalization in the Hippocampal Formation. Cell, 183(5), 1249–1263. https://doi.org/10.1016/j.cell.2020.10.024

Whittington, J. C., McCaffary, D., Bakermans, J. J., & Behrens, T. E. (2022). How to build a cognitive map: insights from models of the hippocampal formation. arXiv. https://doi.org/10.48550/arXiv.2202.01682

Zandbelt, B. (2017) Slice display. figshare. Available at https://doi.org/10.6084/m9.figshare.4742866.

